# A moments-based approach for inferring mechanisms of transcriptional regulation using nascent RNA data

**DOI:** 10.64898/2026.01.03.697452

**Authors:** Kunwen Wen, Yu Liao, Jiandong Wang, Sandeep Choubey, Feng Jiao

## Abstract

Unraveling the kinetics of transcriptional regulation is a central challenge in molecular biology. Traditional inference methods based on cell-to-cell mRNA variability are limited because mRNA levels reflect not only transcriptional activity but also post-transcriptional processes. Recent advances in quantifying nascent RNAs across isogenic cells provide a more direct readout of transcriptional dynamics. Here, we present a moments-based theoretical framework that leverages nascent RNA data to infer transcriptional kinetics. We derive closed-form expressions for the first two time-dependent moments of nascent RNA across three canonical transcription models and demonstrate that nascent RNA exhibits distinct transient overshoots and higher Fano factors compared to mature mRNA. Through analysis of synthetic datasets, we show that these features enable more accurate estimation of transcriptional kinetics and allow reliable model selection, distinguishing single from multi-pathway promoter activation with over 75% accuracy even from small sample sizes. We further validated our model selection method using real experimental transcription datasets in yeast and *E. coli* cells, with nascent RNA demonstrating superior performance in identifying cross-talk pathway regulation. Our framework thus offers a computationally efficient and mechanistically informative approach for decoding transcriptional dynamics, emphasizing the advantages of time-dependent nascent RNA measurements over mature mRNA in capturing the true kinetics of transcriptional regulation.

## Introduction

Single cells tightly regulate the expression of various genes in response to both environmental and intracellular signals. A key step in this regulatory process is transcription, during which transcription factors bind to the promoter regions of genes and modulate their expression. Deciphering the mechanisms underlying transcriptional regulation remains a central challenge in modern biology. The small number of molecules involved makes transcription inherently stochastic, leading to significant cell-to-cell variability within a genetically identical cell population, known as transcriptional noise. Transcriptional noise is observed in both nascent RNA, associated with actively transcribing RNA polymerases, and mature messenger RNA (mRNA) [1–3] (Fig. 1a). Advances in single-molecule imaging and single-cell RNA sequencing have revolutionized our ability to quantify this variability by enabling precise counting of nascent RNA and mRNA molecules at the single-cell level [4, 5]. These measurements have been instrumental in advancing a mechanistic understanding of transcriptional regulation through an effective dialogue between theory and experiment. In particular, theoretical studies propose kinetic models of transcription, whose predictions are systematically tested against data by tuning model parameters to capture observed variability, thereby establishing a framework that links molecular mechanisms of transcription to measurements of nascent RNA and mRNA.

**Figure 1:**
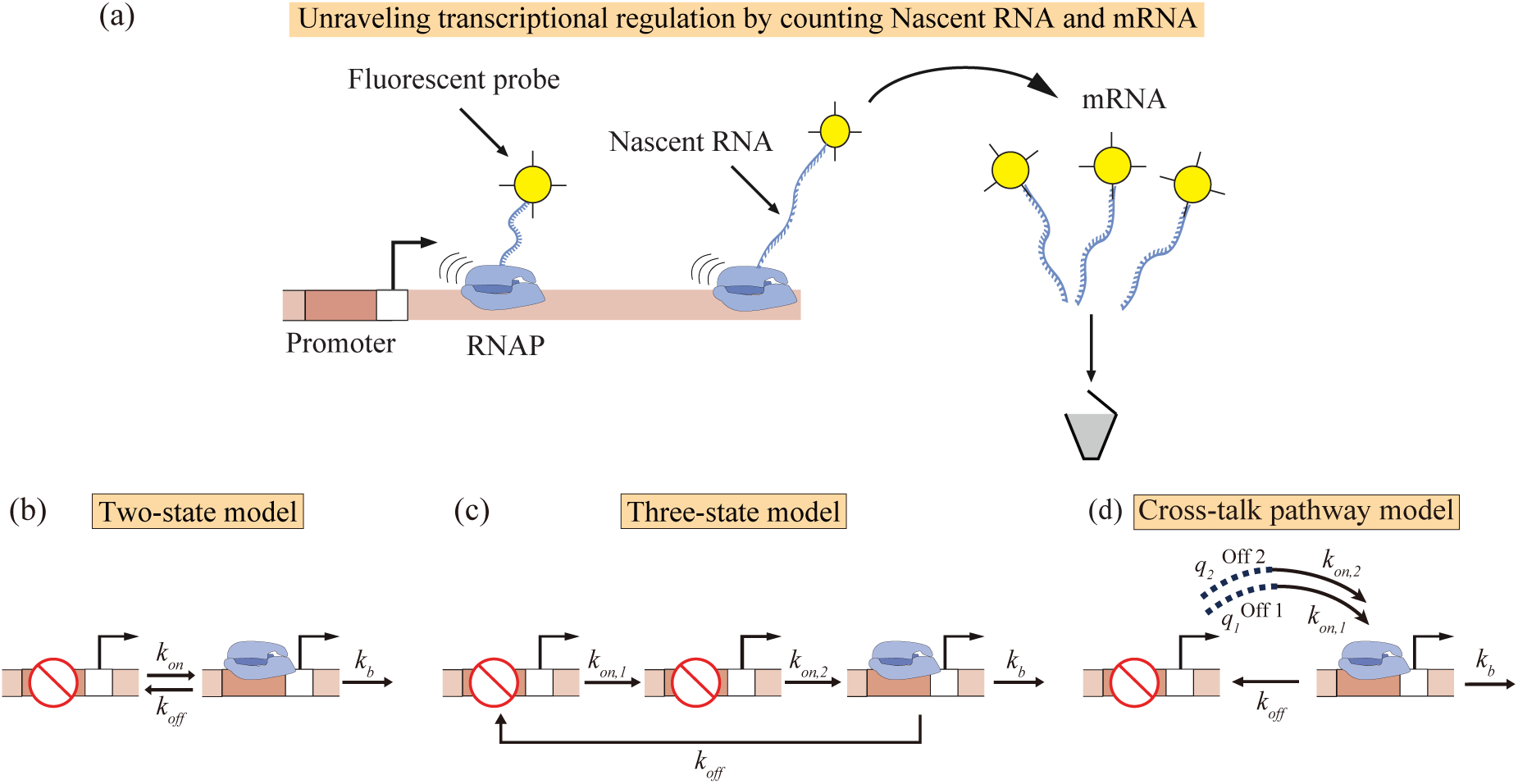
Schematic of different models of transcriptional regulation. **(a)** Following transcriptional initiation, each RNA polymerase (RNAP) molecule traverses a gene (of length *L*) at a speed *v*, generating nascent RNA in the process. The time required to traverse the gene is therefore *T* = *L/v*. Upon completing transcription of the gene, RNAP molecules dissociate from the DNA, thereby yielding mRNA molecules. Subsequently, mRNA undergoes degradation with a first-order rate constant *k*_d_. **(b)** The two-state model describes gene switching between inactive (off) and active (on) states, defined by rates *k*_on_ (activation) and *k*_off_ (inactivation), respectively. In the active state, mRNA or nascent RNA is synthesized at rate *k*_b_. **(c)** The three-state model incorporates ”refractory” behavior: upon exiting the active state at rate *k*_off_, the gene proceeds through two consecutive inactive states with transition rates *k*_on,1_ and *k*_on,2_ before reactivating. **(d)** The cross-talk pathway model involves gene activation via two signaling pathways, with respective rates *k*_on,1_ and *k*_on,2_. Their competitive dynamics are captured by selection probabilities *q*_1_ and *q*_2_ = 1 − *q*_1_.

Kinetic models of transcription have been particularly successful in explaining different regimes of transcriptional noise. For example, housekeeping genes often display Poissonian fluctuations in mRNA levels, which can be captured by a simple birth-death model in which mRNA is synthesized at a constant rate and degraded via first-order decay [2]. In contrast, many genes exhibit super-Poissonian noise, well explained by the two-state (also known as ON-OFF model) [1, 5]. In this model, the promoter stochastically switches between an transcriptionally inactive state and an active state (Fig. 1b). The episodic activation of the promoter gives rise to transcriptional bursts, resulting in variability exceeding that of the Poissonian case. Despite its wide applicability, the predictive power of the two-state model is limited because it neglects important biological mechanisms. In particular, it assumes that the durations of both active and inactive states follow exponential distributions. The exponential assumption for the active and inactive states have been shown to be inconsistent with recent experimental observations. For instance, studies in both mammalian and bacterial cells indicate that inactive periods can deviate significantly from an exponential distribution, instead exhibiting a peaked, non-exponential profile [6–8]. This observation suggests that promoter activation may involve multiple rate-limiting steps, giving rise to refractory behavior. Such dynamics are better described by a three-state model (Fig. 1c), in which the inactive period is subdivided into sequential steps [7, 9]. Mechanistically, this refractory behavior likely arises from the inherently multi-step nature of promoter activation, which requires chromatin remodeling as well as the binding and unbinding of transcription factors or RNA polymerase [10, 11]. Similarly, studies of PHO5 transcription in yeast have revealed peaked distributions duration in active states, indicating the involvement of non-equilibrium mechanisms in transcriptional regulation [12]. These observations suggest that transcriptional regulation often relies on mechanisms more complex than hitherto appreciated.

The advent of high-throughput single-cell data offers a unique opportunity to uncover the complex regulatory mechanisms of transcription through data-driven, backward modeling approaches, in which models are inferred directly from experimental observations. In fact, Bayesian Monte Carlo methods have been used to infer gene expression kinetics and quantify parameter uncertainty from smFISH-based mRNA counts in neurons for the Npas4 gene [13]. Similarly, approximate Bayesian computation has been applied to single-cell transcriptomics data to infer transcriptional burst kinetics using a generalized, non-Markovian telegraph model, yielding more accurate estimates of burst size and frequency than classical Markovian models [14]. In another study, Bayesian inference techniques combined with dual-color MS2/PP7 reporters in Drosophila embryos enabled real-time, single-cell inference of transcription cycle parameters-initiation, elongation, and cleavage-revealing significant variability in elongation rates and a mechanistic link between RNA polymerase density (or equivalently nascent RNA density) and transcript cleavage efficiency [15]. While inference methods using nascent RNA and mRNA distributions offer potential for revealing mechanistic details of transcriptional regulation, the robustness and consistency of models and parameters inferred from these two measurement types remain uncertain. In particular, the impact of the temporal dynamics of nascent RNA and mRNA on model selection and parameter estimation has not yet been thoroughly investigated.

To this end, we developed a moments-based inference approach that uses nascent RNA and mRNA data. By deriving analytical expressions for the first and second moments of nascent RNA distributions for three canonical models of transcription, we identified distinct dynamical features of nascent RNA, including a pronounced overshoot in the mean and elevated Fano factors, which preserve transcriptional regulatory signatures more reliably than mRNA. This framework enables accurate fitting of time-dependent nascent RNA data and outperforms mRNA-based methods in both parameter estimation-especially improving the inference of gene inactivation rates-and model selection, achieving over 75 percent accuracy in distinguishing single-versus multi-pathway activation mechanisms, even with limited data. These findings highlight the advantages of leveraging nascent RNA’s dynamical properties for inferring mechanistic details of transcriptional regulation.

## Results

### Time-Dependent analytical expressions for the first two moments of Nascent RNA distributions for different models of transcriptional regulation

To systematically compare moments-based approaches for inferring transcriptional regulatory models from nascent RNA and mRNA data, we analyze three distinct models of transcriptional regulation. In this section, we consider three models of transcriptional regulation: i) Two-state model, ii) Three-state model, and iii) Cross-talk pathway model. Our goal is to compute time-dependent analytical expressions for the first two moments of the nascent RNA and mRNA distributions for these models. To compute the moments of nascent RNA distributions, we map these models to a queueing process. Since nascent RNAs are produced when the gene is in an active state for all the models we consider, the time intervals between successive initiation events follow independent and identically distributed distributions-resembling a renewal process. Hence, our models can be described using the well-studied *G/D/*∞ system from queueing theory [16]. In this framework, *G* denotes a general waiting time distribution, *D* a deterministic service time, and ∞ an infinite number of servers. Leveraging this mapping, we derive analytical formulas for the mean and second moment of the nascent RNA distributions for the different model. Moreover, we use a master equation-based approach to obtain the time-dependent moments of mRNA distribution [9, 17, 18]. Below, we discuss the results in detail.

### Two-state model

First, we consider the two-state model of transcriptional regulation [1, 19, 20], as illustrated in Fig. 1b. The two-state model of transcriptional regulation has been remarkably successful in explaining transcriptional bursting across diverse systems. In mammalian cells, single-molecule FISH studies have shown that many genes produce highly variable mRNA copy numbers across an identical cell population, a phenomenon that is well explained by the two-state model of gene expression [21]. Similarly, more recent live-imaging studies in Drosophila embryos have shown that several developmental genes undergo bursty transcription, a behavior that is consistent with the two-state model [22]. The two-state model posits that a gene transitions stochastically between an inactive state and an active state, governed by activation rate *k*_on_ and inactivation rate *k*_off_. Nascent RNA molecules are synthesized at a rate *k*_b_ when the gene is in the active state. After transcribing a gene, RNA polymerase molecules detach from the DNA, leading to the production of mature messenger RNA molecules. Given the short lifetime of nascent RNA compared to mRNA, we model the production processes of both nascent RNA and mRNA as a single step for mRNA synthesis [10, 19]. For mRNA molecules, their degradation, which includes both decay and dilution during cell division, is characterized by a decay rate *k*_d_. In contrast, the residence time of nascent RNA on a gene, i.e., the time it takes for an RNA polymerase to transcribe a gene, is modeled as a deterministic process. The time it takes for an RNAP molecules to transcribe a gene is *T* = *L/v*, where *L* denotes the gene length and *v* represents the speed of elongation along the gene[6, 20, 23].

We compute the transient mean *M*_1_(*t*) and the second moment *S*_1_(*t*) for nascent RNA, along with the corresponding transient mean *M*_2_(*t*) and the second moment *S*_2_(*t*) for mRNA, as summarized in Table 1 (see Supplementary Material for details).

**Table 1:** Exact formulas for the transient mean *M*_1_(*t*) and second moment *S*_1_(*t*) of the nascent RNA distribution [23], as well as the mean *M*_2_(*t*) and second moment *S*_2_(*t*) of the mRNA distribution [17] in the two-state model.

| | $t \leq T$ | $t > T$ | Steady state value |
| --- | --- | --- | --- |
| $M_1(t)$ | $a + bt - ae^{-ct}$ | $bT + ae^{-ct} (e^{Tc} - 1)$ | $bT$ |
| $S_1(t)$ | $M_1(t) + b^2T^2 + 4abt - 2a^2ct e^{-ct} + (2a^2 - 4ab/c)(1 - e^{-ct})$ | $M_1(t) + b^2T^2 + 2a^2e^{-ct}(e^{cT} - cT - 1) + (ab/c)(e^{-ct} + e^{-cT})[1 + e^{cT}(cT - 1)]$ | $bT(bT + 1) + \frac{ab(e^{-cT} + cT - 1)}{c}$ |
| $M_2(t)$ | $\frac{b}{k_d} - \frac{b}{(k_d - c)}e^{-ct} + \frac{bc}{k_d(k_d - c)}e^{-k_d t}$ | | $\frac{b}{k_d}$ |
| $S_2(t)$ | $\frac{b[k_d(k_d + c) + k_b(k_d + k_{\text{on}})]}{k_d^2(k_d + c)} - \frac{b[2k_d^2 - k_dc + 2k_b(k_d - k_{\text{off}})]}{k_d(k_d - c)(2k_d - c)}e^{-ct} - \frac{bc k_d + 2bk_{\text{on}}k_b}{k_d^2(c - k_d)}e^{-k_d t} + \frac{k_{\text{on}}k_b^2(k_{\text{on}} - k_d)}{k_d^2(c - 2k_d)(c - k_d)}e^{-2k_d t} + \frac{2bk_b k_{\text{off}}}{k_d(k_d + c)(k_d - c)}e^{-(k_d + c)t}$ | | $\frac{b[k_d(k_d + c) + k_b(k_d + k_{\text{on}})]}{k_d^2(k_d + c)}$ |
| Coefficients | $c = k_{\text{on}} + k_{\text{off}}, a = k_b k_{\text{off}}/c^2, b = k_b k_{\text{on}}/c.$ | | |

### Three-state model and cross-talk pathway model

Second, we consider the three-state model of transcriptional regulation, as shown in Fig. 1c: this model assumes the promoter has three possible states (one active and two inactive), where the inactive period involves two rate-limiting steps characterized by activation rates *k*_on,1_ and *k*_on,2_, and exhibits a “refractory” behavior-i.e., the gene transitions through two consecutive inactive states prior to reactivation [7, 9]. Notably, recent experiments in mammalian and bacterial cells indicate that the inactive periods of some genes may exhibit a non-exponential peaked distribution [6–8], a feature well-captured by the three-state model [7, 9]. The non-exponential nature of the inactive period is attributed to a multi-step biochemical process at the promoter, which includes chromatin remodeling along with the binding and unbinding dynamics of transcription factors and RNA polymerases [10, 11]. Third, we consider the cross-talk pathway model [18, 24]. Both the two-state and three-state models involve only a single gene activation pathway. However, gene activation is often regulated through multiple signaling pathways-this is captured by the cross-talk pathway model (Fig. 1d). This model describes two transcription factors or enhancers competing to bind the promoter: one activates the gene via a weak signaling pathway with rate *k*_on,1_, and the other via a strong pathway with a higher rate *k*_on,2_ *> k*_on,1_. The probabilities of activation through the weak and strong pathways are *q*_1_ and *q*_2_ with *q*_1_ + *q*_2_ = 1, respectively. Recent studies have shown that the cross-talk model accurately reproduces the rapid overshooting of mRNA levels observed in experiments: for example, in mouse fibroblasts induced by tumor necrosis factor [24, 25], and in *E. coli* induced by arabinose or isopropylthio-L-D-galactoside [26]. Such overshooting cannot be explained by the two-state or three-state models, which produce nearly monotonic increases in mRNA mean level [9, 24].

In the three-state model and the cross-talk pathway model, the transient mean *M*_1_(*t*) and the second moment *S*_1_(*t*) for nascent RNA, the transient mean *M*_2_(*t*) and the second moment *S*_2_(*t*) for mRNA, are separately given in Table 2 (see detailed calculations in the Supplementary Material).

**Table 2:**
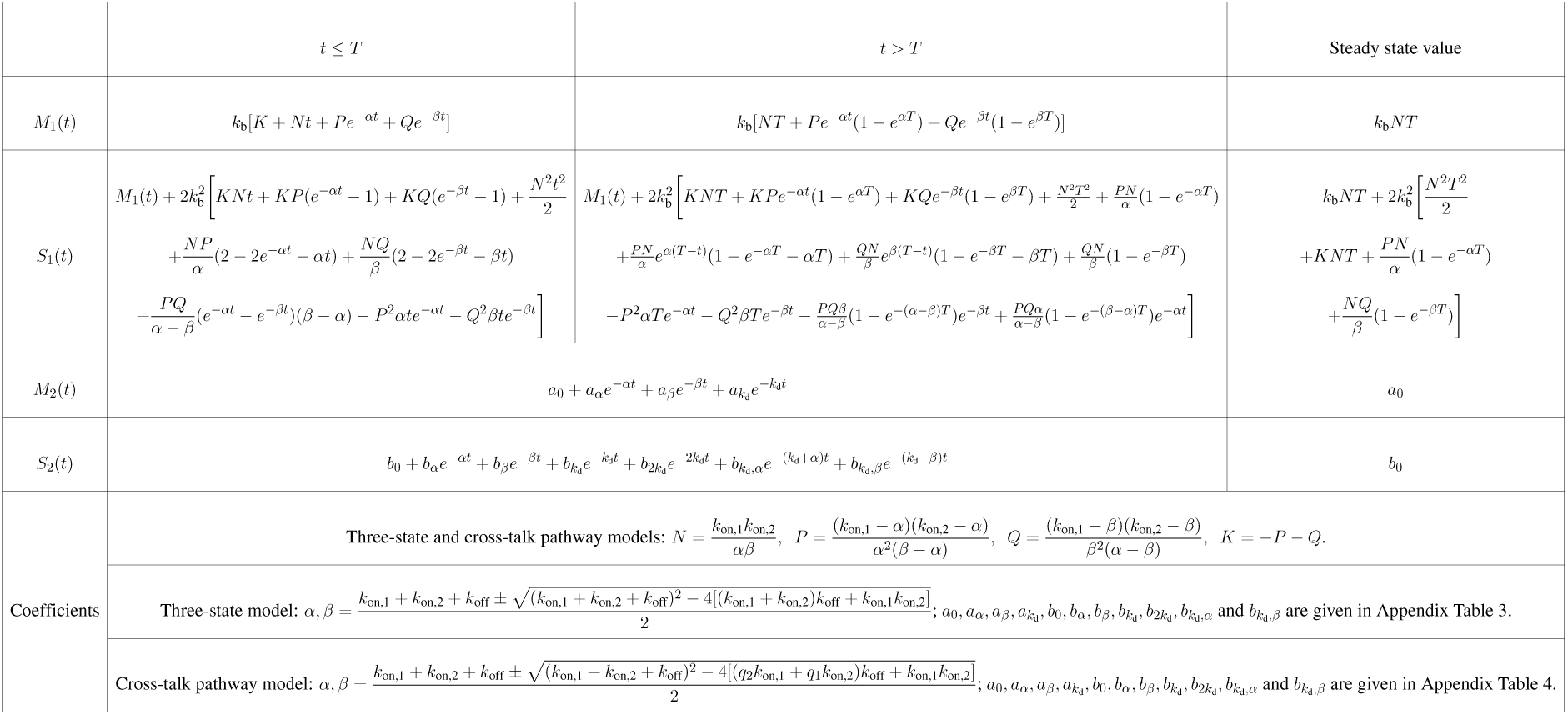
Exact formulas of the three-state and cross-talk pathway models for transient mean *M*_1_(*t*) and the second moment *S*_1_(*t*) of nascent RNA distribution (see detailed calculations in the Supplementary Material), and mean *M*_2_(*t*) and the second moment *S*_2_(*t*) of mRNA distribution [9, 18].

The exact formulas for mean and second moment for the three models were further verified using Gillespie stochastic simulation algorithm (SSA) with 10^4^ cell samples (Supplementary Fig. S1) [27]. The first two moments of nascent RNA distribution for the three models can be written in terms of the products *k*_on_*T, k*_off_*T, k*_on,1_*T, k*_on,2_*T, k*_b_*T* , as well as the ratio *t/T* . Similarly, the exact expressions for the first two moments of the mRNA distribution depend on the ratios *k*_on_*/k*_d_*, k*_off_*/k*_d_*, k*_on,1_*/k*_d_*, k*_on,2_*/k*_d_*, k*_b_*/k*_d_, and the product *k*_d_*t*. To streamline the calculations and improve clarity, we set *T* = 1 and *k*_d_ = 1 for nascent RNA and mRNA, respectively [23, 26]. This normalization ensures that nascent RNA and mRNA exhibit the same steady-state mean level under identical system parameters, which facilitates the comparison of analytical methods using synthetic data for the two types of RNA. Consequently, all model parameters presented henceforth (with the exception of *q*_1_ and *q*_2_ in the cross-talk pathway model) should be interpreted as follows: For nascent RNA, each parameter represents the product of its actual value and *T* ; For mRNA, each parameter represents the ratio of its actual value to *k*_d_. Likewise, the time variable *t* is normalized to align time scales across models: For nascent RNA, *t* corresponds to real time divided by *T* ; For mRNA, *t* corresponds to real time multiplied by *k*_d_.

### Dynamics of the mean and Fano factor of nascent RNA and mRNA

With the exact formulas for the mean, variance, and the Fano factor (ratio of variance and mean) of the nascent RNA and mRNA distributions for the three models at hand, we can quantitatively assess the dynamical differences between them. To this end, we randomly select 100 parameter sets (*b_f_ , k*_off_*, k*_b_) where *b_f_* and *k*_off_ are within the range (0.1, 3) and *k*_b_ is in (5, 30) (Supplementary Table S1). Here, *b_f_* refers to the transcriptional burst frequency, which is defined as follows: *b_f_* = *k*_on_ for the two-state model; *b_f_* = 1*/k*_on,1_ + 1*/k*_on,2_ for the three-state model; and *b_f_* = 1*/*(*q*_1_*/k*_on,1_ + *q*_2_*/k*_on,2_) for the cross-talk pathway model [9, 24], respectively. For both mRNA and nascent RNA in all three models, the exact formula for the steady-state mean *M* ^∗^ is expressed as *M* ^∗^ = *k*_b_*b_f_ /*(*b_f_*+ *k*_off_). Given a parameter set (*b_f_ , k*_off_*, k*_b_), we set the other parameters of each model as follows to maintain the same mean level across all models: *k*_on_ = *b_f_* for the two-state model; *k*_on,1_ = *k*_on,2_ = 2*/b_f_* for the three-state model; and for the cross-talk pathway model, *k*_on,1_ ≡ 0.1, and *q*_2_ = (1*/k*_on,1_ − 1*/b_f_* )*/*(1*/k*_on,1_ − 1*/k*_on,2_) by reversing the expression of *b_f_* [28]. To ensure *q*_2_ ∈ (0, 1), *k*_on,2_ must be greater than *b_f_* . Thus, we randomly selected values of *k*_on,2_ within the range (*b_f_ ,* 8).

We then plot the mean and Fano factor of mRNA and nascent RNA asgainst time *t* for the three models for each of the 100 randomly selected parameter sets (*b_f_ , k*_off_*, k*_b_). As shown in Fig. 2a, the dynamics of the mean nascent RNA level exhibit a more pronounced “overshoot” compared to those of mRNA; the mean nascent RNA level rises very rapidly in the initial time period, reaches a distinct peak, and then quickly decreases to approach its steady-state value. Interestingly, the mean level of nascent RNA is consistently higher than that of mRNA. Furthermore, the cross-talk pathway model yields significantly higher peaks in the mean levels compared to the two-state and three-state models. As shown in Fig. 2b, the Fano factor of nascent RNA initially increases more slowly than that of mRNA, but subsequently accelerates, eventually surpassing the mRNA values. As a consequence, nascent RNA distribution exhibits higher values of Fano factor than mRNA in most cases.

**Figure 2:**
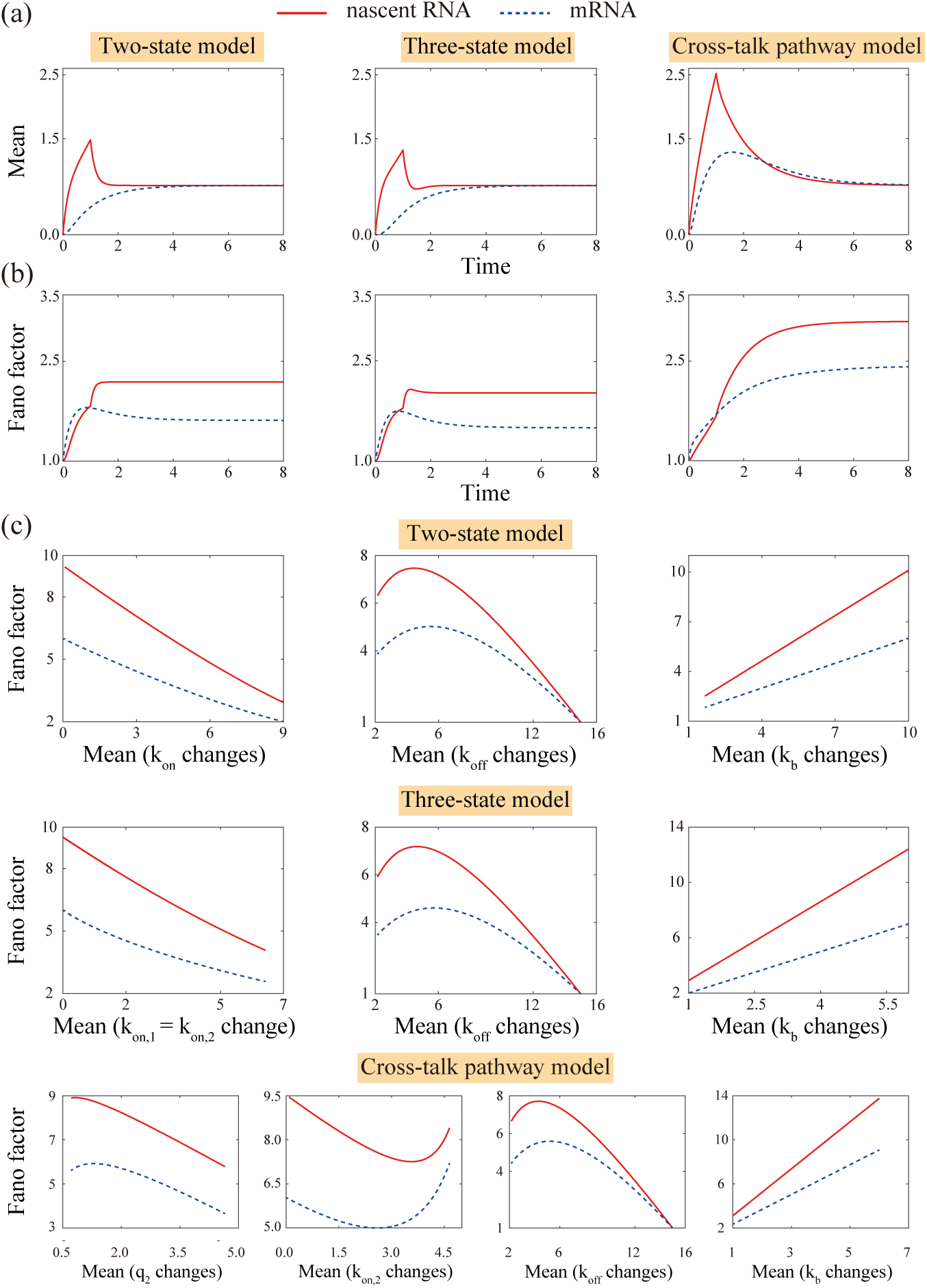
Dynamic characteristics and steady-state correlations of nascent RNA and mRNA. **(a)** Time courses of the mean *M* (*t*) for nascent RNA and mRNA under the two-state, three-state, and cross-talk pathway models. nascent RNA displays a more pronounced “overshoot” behavior: it rises rapidly, showd a peak, and approaches the steady-state value *M* ^∗^, whereas mRNA lacks such a distinct overshoot. Among these models, the cross-talk pathway model produces significantly higher *M* (*t*) peaks (exceeding three times *M* ^∗^). **(b)** Time courses of the Fano factor *η*(*t*) for nascent RNA and mRNA across the three models. nascent RNA *η*(*t*) increases more slowly than mRNA *η*(*t*) in the initial phase but accelerates afterward, eventually surpassing and remaining higher than mRNA *η*(*t*) for most of the time course. **(c)** Plots of the steady-state Fano factor *η*^∗^ versus the steady-state mean *M* ^∗^ for nascent RNA and mRNA across the three models (generated by varying one parameter at a time while fixing others). Curves exhibit distinct monotonic behaviors (decreasing, increasing, or non-monotonic). Nascent RNA and mRNA share the same monotonic trends for a given parameter variation, but nascent RNA consistently shows higher *η*^∗^ values than mRNA at the same *M* ^∗^.

Next, we examine how the Fano factor depends on the mean in the steady state. For each of the three models, we vary one parameter while keeping all other parameters fixed, which causes corresponding changes in mean and Fano factor. We then plot the Fano factor as a function of the mean. As shown in Fig. 2c, the Fano factor exhibits distinct monotonic behaviors with respect to the mean when a single model parameter is varied. We note that when a single parameter is tuned, the Fano factor behaves similarly for both nascent RNA and mRNA. However, for identical mean values, the Fano factor of nascent RNA is consistently higher than that of mRNA.

Our results show that the coupling of stochastic promoter state switching with a deterministic RNA degradation process, as in the case of nascent RNA, generates higher transcriptional noise than its coupling with a stochastic RNA degradation process, as in the case of mRNA. This finding carries important implications for model inference, as we delineate in the ensuing sections.

### Parameter estimation using data for the first two moments of nascent RNA and mRNA distributions

To assess the reliability of parameter estimation based on the moments of nascent RNA and mRNA distributions, we use SSA to generate synthetic data for the first two moments of both species. This is done for each of the three models, across 100 randomly sampled parameter sets (*b_f_ , k_f_ , k_b_*), while varying the number of sampled cells *N* and the number of time points *n*. In other words, we sample *N* values for the first two moments and the Fano factor at time points *t* = *t̄/n,* 2*t̄/n,* · · · *, nt̄/n*, where *t̄* represents the time at which the system reaches steady state (see Supplementary Materials). In this study, we considered 4 cell samples with *N* = 10^2^, 10^3^, 5 × 10^3^, 10^4^ and 5 time point numbers *n* = 2, 4, 8, 12, 24. These values cover the ranges of cell numbers and time point numbers observed in experimental data measured using single-cell RNA sequencing [3, 29, 30].

For each synthetic dataset of the moments of nascent RNA and mRNA for different sample sizes, the traditional approach to parameter estimation involves using the exact formulas of RNA mean *M* (*t*) and second moment *S*(*t*) to fit the synthetic data by minimizing the sum-of-squares differences *D*(*θ*) (i.e., least-squares fitting) [25, 29, 31, 32]

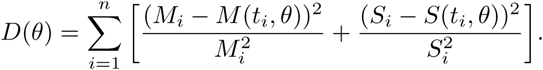

Here, *θ* denotes the model parameter set; *n* represents the number of measurement time points; *M_i_* and *S_i_* are the synthetic data of the mean and the second moment at time *t_i_ >* 0, respectively; and *M* (*t_i_, θ*) and *S*(*t_i_, θ*) are the simulated *M* (*t*) and *S*(*t*) at *t* = *t_i_*, obtained using their exact formulas, respectively. Each minimization optimization was executed 100 times with randomly generated initial parameters.

Simultaneously fitting of RNA *M* (*t*) and *S*(*t*) data enables rapid parameter estimation, thanks to the efficient computation of their exact formulas. To verify the reliability of this estimation method, we generated synthetic data using the two-state model. As shown in Fig. 3a-where the synthetic data was generated with 4 time points and 10^3^ cell samples-there is a relatively good match between the fitted curves and the synthetic data for the first two moments. However, non-negligible discrepancies are observed between the fitted curves and the synthetic data for the Fano factor as a function of time. To achieve higher fitting accuracy, we incorporated the Fano factor *η*(*t*) into the cost function; we minimized the corrected sum-of-squares difference *D_c_*(*θ*):

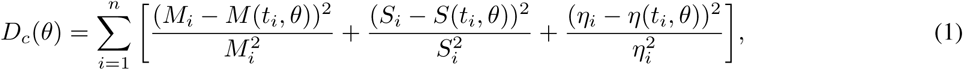

where *η_i_*represents the Fano factor of nascent RNA and mRNA, as computed from synthetic data at time *t_i_ >* 0, and *η*(*t_i_, θ*) denotes simulated Fano factor values at *t* = *t_i_* , derived from its exact formulas. As illustrated in Fig. 3a, minimizing the modified *D_c_*(*θ*) improves fitting accuracy of the Fano factor of nascent RNA data by more than ten-fold compared to the previous case. This improvement is quantified using the average absolute distance (AAD) between the fitted curves and the synthetic nascent RNA data, where AAD is defined as ^L^*^n^* |*A_i_* − *A*(*t_i_*)|*/n*, where *A_i_* is synthetic data, and *A*(*t_i_*) are values of the fitted curve at time *t_i_*. Additionally, compared to fitted curves of mean and second moment, obtained by minimizing *D*(*θ*), those obtained by minimizing *D_c_*(*θ*) are more effective at capturing the true profiles of *M* (*t*) and *S*(*t*), both near their peaks and in the post-peak regions.

**Figure 3:**
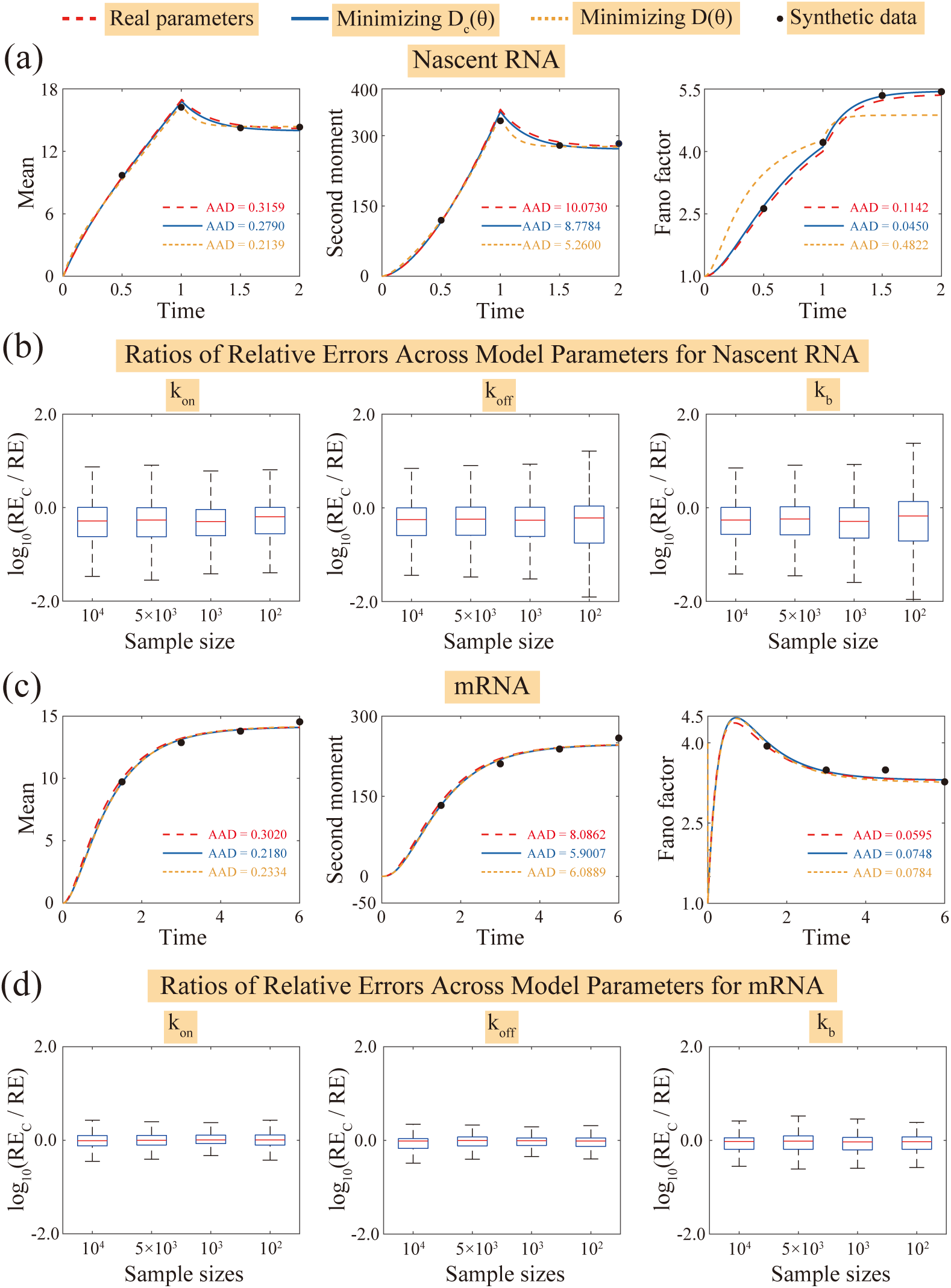
Performance of moment-based parameter estimation methods for nascent RNA and mRNA synthetic data. **(a)** Fitting of nascent RNA data via minimization of the corrected sum-of-squares difference *D_c_*(*θ*) shows a significantly better match with the synthetic data compared to fitting via minimization of *D*(*θ*), with an over 10-fold improvement for fitting Fano factor data in the average absolute distance (AAD). **(b)** Ratios of relative errors for each parameter (*k*_on_*, k*_off_*, k*_b_) estimated via minimization *D_c_*(*θ*) versus *D*(*θ*) when fitting nascent RNA data. Results are derived from 100 parameter sets, 4 cell sample sizes (10^2^, 10^3^, 5 × 10^3^, 10^4^), and 5 time point numbers (2, 4, 8, 12, 24). Over 70% of these ratios are *<* 1 (or log_10_(ratios) *<* 0) across all parameters, confirming that estimates from *D_c_*(*θ*) are more accurate than those from *D*(*θ*). **(c)** Fitting of mRNA data via minimization of *D_c_*(*θ*) and *D*(*θ*) yields comparable performance, owing to the lack of sharp dynamic peaks in mRNA profiles. **(d)** For mRNA data, under the same conditions as in (b), ratios of relative errors for each parameter are estimated via minimization of *D_c_*(*θ*) versus *D*(*θ*) during fitting. Minimization of *D_c_*(*θ*) yields performance comparable to that of *D*(*θ*) in estimating *k*_on_, while showing a slight advantage in estimating *k*_off_ and *k*_b_. This is consistent with the modest improvements in mRNA data fitting observed in (c).

To further verify whether the moment-based fitting method by minimizing *D_c_*(*θ*) as defined in (1)-can yield more reliable parameter estimates than minimizing *D*(*θ*), we analyzed nascent RNA synthetic data generated via the two-state model. In particular, we defined the relative errors RE*_c_*(*b_f_* ), RE*_c_*(*k*_off_) and RE*_c_*(*k*_b_) as the discrepancies between the estimated parameters and their true values (*b_f_ , k*_off_*, k*_b_), when we minimize *D_c_*(*θ*). Correspondingly, we defined RE(*b_f_* ), RE(*k*_off_), and RE(*k*_b_) for inferred parameter values by minimizing *D*(*θ*). Here, the relative error is defined as |*P̄* − *P* |*/P* , where *P* denotes the true parameter value, and *P̄* denotes the inferred parameter value. We then computed the ratios, RE*_c_*(*b_f_* )/RE(*b_f_* ), RE*_c_*(*k*_off_)/RE(*k*_off_), and RE*_c_*(*k*_b_)/RE(*k*_b_) for the different parameters. These computations were carried out for 100 distinct parameter sets (*b_f_ , k*_off_*, k*_b_), 5 different numbers of measured time points (*n* = 2, 4, 8, 12, 24), and 4 cell sample sizes (*N* = 10^2^, 10^3^, 5 × 10^3^, 10^4^). For each of the 3 parameters and each of the 4 sample sizes, a total of 100 × 5 = 500 ratio values were generated. These results are visualized in Fig. 3b and Supplementary Fig. S2. As shown, over 70% of the ratio values are less than 1 for every parameter and every sample size. This indicates that parameter estimates derived by minimizing *D_c_*(*θ*) are more accurate than those obtained by minimizing *D*(*θ*).

We further validated parameter estimation using synthetic mRNA data generated by the two-state model. Notably, the two-state model does not produce sharp dynamic peaks in mean and the second moments of mRNA as functions of time. This crucial property ensures that incorporating the Fano factor into the cost function does not affect the inference procedure; in other words, parameter estimation remains unchanged whether we use *D*(*θ*) or *D_c_*(*θ*), as illustrated in Fig. 3c. This became evident when we calculated the ratios RE*_c_*(*b_f_* )/RE(*b_f_* ), RE*_c_*(*k*_off_)/RE(*k*_off_) and RE*_c_*(*k*_b_)/RE(*k*_b_) using the mRNA synthetic data, under the same parameter sets, sample sizes, and numbers of time points as applied in Fig.3b. The results indicate that minimizing *D_c_*(*θ*) and *D*(*θ*) yields similar performance in estimating *b_f_* , while minimizing *D_c_*(*θ*) performs slightly better than minimizing *D*(*θ*) in estimating *k*_off_ and *k*_b_, see Fig. 3d and Supplementary Fig. S3.

Overall, our computational analysis demonstrates that incorporating the Fano factor into the parameter estimation process substantially improves the accuracy of model fitting for nascent RNA data compared to using only the first two moments. In contrast, for mRNA data, the improvement is minimal.

### Reliability of parameter estimation with nascent RNA data

To further explore the reliability of parameter inference using nascent RNA data, we tune the cell sample size *N* and number of time points *n*. Additionally, we examine whether nascent RNA data or mRNA data yields more reliable parameter estimates. A parameter is considered accurately estimated if the absolute relative error between the estimated and true parameters does not exceed 0.2 [26, 31]. For each fixed pair (*N, n*) across 100 parameter sets (*b_f_ , k*_off_*, k*_b_), a parameter’s estimation is deemed reliable if it is accurately estimated for at least 70 of the tested parameter sets, i.e., the accuracy of estimation exceeds 70%.

We first analyzed synthetic nascent RNA data generated using the two-state model, fitting it by minimizing *D_c_*(*θ*) to estimate the parameters *b_f_* = *k*_on_*, k*_off_, and *k*_b_. As shown in the first row of Fig. 4a, for the smallest sample size (*N* = 100), the estimate of *k_b_* is the most reliable, whereas the estimates of *b_f_* and *k_f_* are the least reliable. For reliable estimation of *k*_b_: a sample size of *N* = 10^2^ requires at least *n* ≥ 4; a sample size of *N* ≥ 10^3^ requires only *n* ≥ 2. For reliable estimation of *b_f_* and *k*_off_: *N* = 10^2^ requires at least *n* ≥ 24; *N* = 10^3^ requires *n* ≥ 4; *N* ≥ 5 × 10^3^ requires *n* ≥ 2. We next examined synthetic mRNA data generated by the two-state model (see the first row of Fig. 4b). Reliable estimation of *k*_b_ requires similar sample sizes and time points, as observed in the fitting of nascent RNA data. The estimated *k*_off_ is the least reliable, demanding at least *N* ≥ 10^3^ and *n* ≥ 4. In summary, to ensure reliable estimation of all parameters (*b_f_ , k*_off_*, k*_b_) of the two-state model: fitting nascent RNA moment data requires either *N* = 10^2^ with *n* ≥ 24, or *N* ≥ 10^3^ with *n* ≥ 4; fitting mRNA data is not feasible with *N* = 10^2^, as it requires at least *N* = 10^3^ and *n* = 4. Thus, fitting nascent RNA moment data may yield more reliable parameter estimation for the two-state model than fitting mRNA moment data.

**Figure 4:**
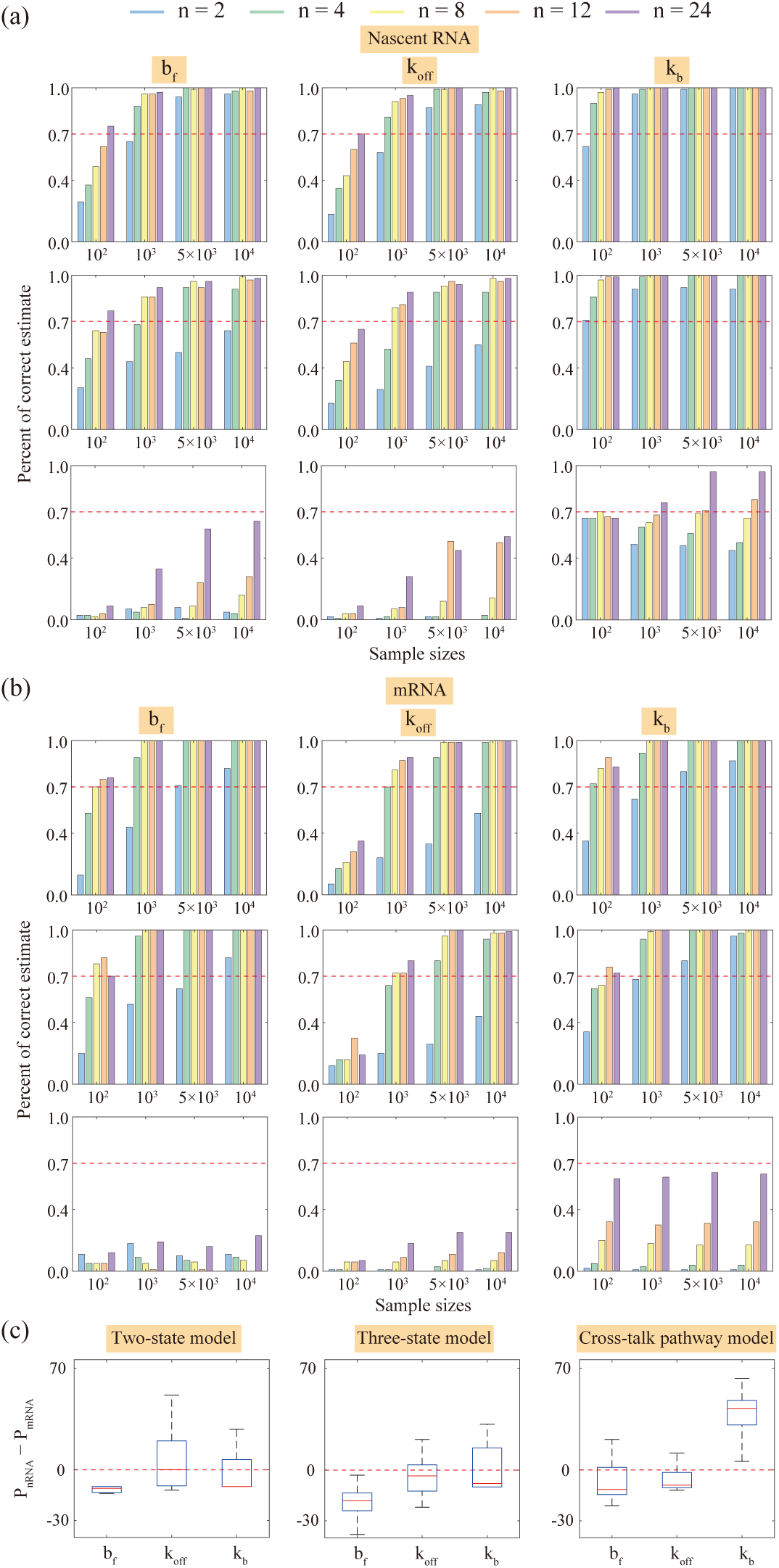
Impact of data type, sample size *N* , and the number of time points *n* on reliability of parameter estimation. **(a)-(b)** Reliability of parameter estimates (*b_f_ , k*_off_*, k*_b_)-from synthetic nascent RNA data in (a) and mRNA data in (b)-as a function of *N* (10^2^, 10^3^, 5 × 10^3^, 10^4^) and *n* (2, 4, 8, 12, 24), for the two-state (1st row), three-state (2nd row), and cross-talk pathway (3rd row) models. A parameter estimates is reliable if ≥ 70% of 100 parameter sets yield accurate results (relative error ≤ 0.2; red horizontal dashed line). For the two-state model, nascent RNA data supports reliable estimation with smaller *N* and *n* compared to mRNA data; for the three-state model, nascent RNA data shows slightly higher reliability for *k*_off_ and *k*_b_ estimates; for the cross-talk model, nascent RNA data yields clearly more reliable estimates than mRNA data. Notably, neither data type achieves ≥ 70% accuracy for *b_f_* and *k*_off_ under tested *N* and *n*, indicating limitations of using only the first two moments for this model. **(c)** Differences in accurate estimation percentages *P*_nascentRNA_ − *P*_mRNA_ for each parameter across all three models. Positive values indicate nascent RNA outperforms mRNA. Nascent RNA consistently outperforms mRNA in *k*_off_ and *k*_b_ estimation across all models, and in *b_f_* estimation for the cross-talk pathway model.

To further validate our conclusion that parameter estimation is more reliable when fitting nascent RNA data compared to mRNA data, we generated both synthetic nascent RNA and mRNA data for the first two moments using the three-state model and cross-talk pathway model. These data were generated under 100 randomly selected parameter sets (*b_f_ , k*_off_*, k*_b_), with varying sample sizes *N* = 10^2^, 10^3^, 5 × 10^3^, 10^4^ and numbers of time points *n* = 2, 4, 8, 12, 24. For the three-state model (see the second rows of Fig. 4a for nascent RNA and Fig. 4b for mRNA), while fitting nascent RNA data, *k*_b_ exhibits the highest estimation reliability, whereas *k*_off_ shows the lowest. To ensure reliable estimation of *k*_off_, at least *N* = 10^3^ with *n* ≥ 8, or *N* ≥ 5 × 10^3^ with *n* ≥ 4 is required. In contrast, the reliability for estimating *k*_off_ is slightly less while fitting mRNA data. For the cross-talk pathway model (see the third rows of Fig. 4a for nascent RNA and Fig. 4b for mRNA), parameter estimation by fitting nascent RNA data is clearly more reliable than by fitting mRNA data. However, the tested cell sample sizes and numbers of time points were insufficient to achieve an accuracy of greater than 70% for both estimated *b_f_* and *k*_off_. This suggests that the data for the first two moments alone do not lead to reliable parameter estimation for the cross-talk pathway model.

To further illustrate the advantage of using nascent RNA data for parameter estimation, we define *P*_nascent RNA_ and *P*_mRNA_ as the percentages of accurate parameter estimates obtained from nascent RNA and mRNA data, respectively. We then calculated the difference *P*_nascentRNA_ − *P*_mRNA_ for each of the following parameters: *b_f_ , k*_off_ and *k*_b_ across the two-state, three-state, and cross-talk pathway models (see Fig. 4c). If *P*_nascentRNA_ − *P*_nascentRNA_ *>* 0, this indicates that using nascent RNA data yields better performance than using mRNA data, and vice versa. For the estimation of *b_f_* : In the two-state and three-state models (which involve a single gene activation pathway), nascent RNA data exhibits slightly lower reliability compared to mRNA data. In contrast, nascent RNA data outperforms mRNA data in the cross-talk pathway model. Furthermore, nascent RNA data consistently shows significantly higher reliability than mRNA data when estimating both *k*_off_ and *k*_b_. Notably, *k*_off_ is the most poorly estimated parameter across models when fitting mRNA data (Fig. 4a,b) [19, 26, 33]. Therefore, using nascent RNA data can significantly improve the accuracy of *k*_off_ estimation, thereby enabling more reliable parameter estimation for the models overall.

### Pathway selection for gene activation: Single vs. multiple pathways using nascent RNA moment data

In the two-state and three-state models, gene activation is regulated by a single pathway, whereas the cross-talk pathway model utilizes two competitive pathways to govern gene activation. To distinguish whether a system is regulated by a single pathway or multiple pathways, a traditional approach involves fitting time-resolved RNA distribution data using the maximum-likelihood method [26, 30]. This process generates a likelihood function, which combined with penalty terms that vary with the number of parameters and sample size, yields different criteria for selecting the optimal model to explain the data. However, this method requires significant computational resources and performs poorly when the sample size and time point numbers are small [26, 30, 31].

We investigated whether nascent RNA moment data can reliably distinguish between data generated by a single activation pathway and that generated by cross-talk pathway. Given a dataset for the mean and Fano factor, we fit the data using the two-state model. If the data is generated by the cross-talk pathway model, the estimated *k*_on_ of the two-state model must be extremely large-this is because its fitting curve must align with the rapid overshoot dynamics of the data (see Fig. 5a). Furthermore, a large *k*_on_ leads to a small Fano factor, as indicated by its steady-state exact formula [1]. This causes the Fano factor fitting curve to deviate significantly from the corresponding data. The model selection procedure is as follows: First, fit the nascent RNA data using the two-state model by minimizing *D_c_*(*θ*) described in (1). Then, calculate the average relative errors (ARE) between the fitting curve and the Fano factor *η*(*t*) data, where ARE is defined as 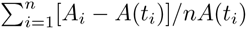, where *A_i_* is synthetic data, and *A*(*t_i_*) are values of the fitted curve at time *t_i_*. If ARE *>* 1 (where 1 serves as the threshold), we select the cross-talk pathway model to explain the data; if ARE ≤ 1, the data are explained by a single activation pathway (e.g., the two-state model or three-state model).

**Figure 5:**
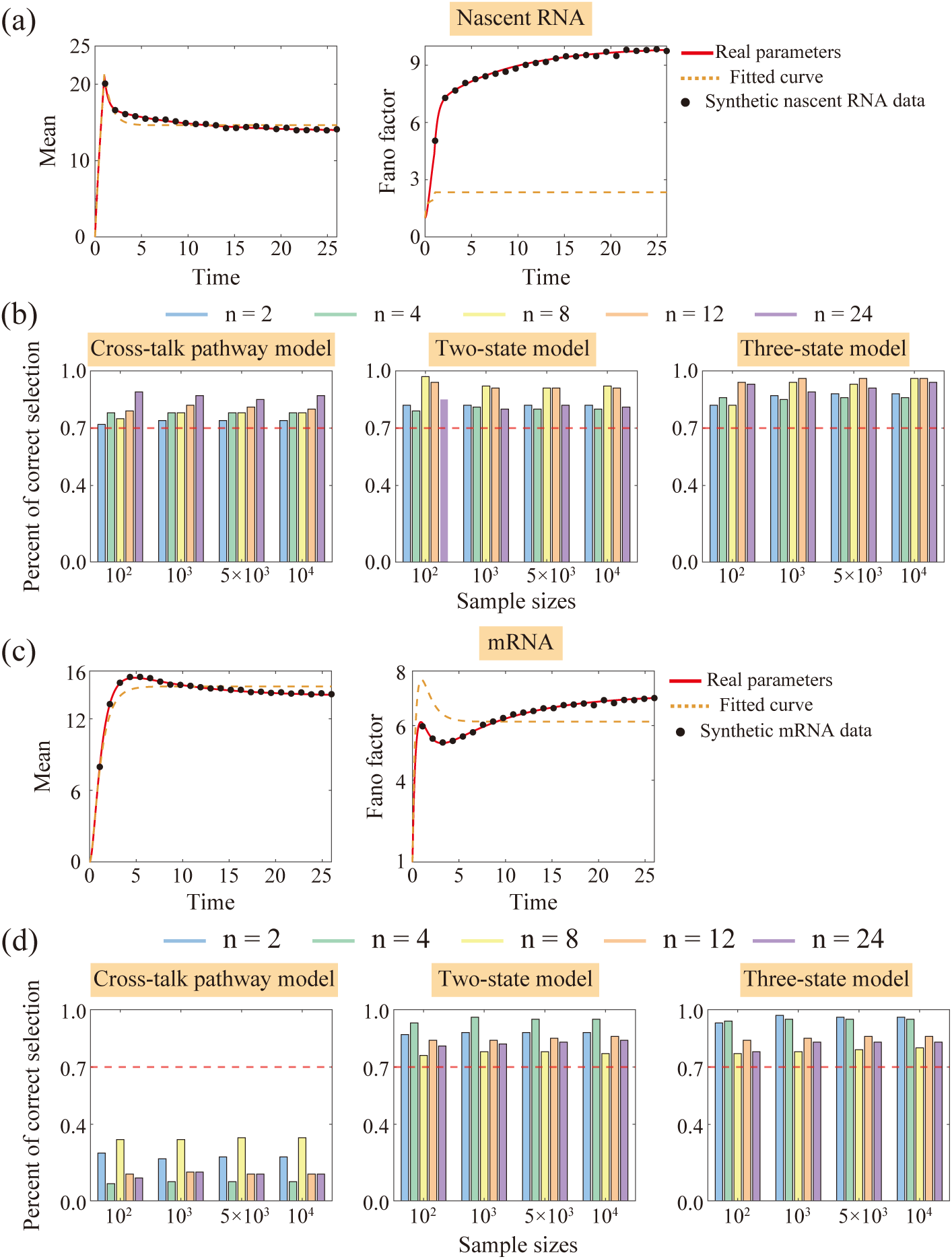
Performance of moment-based model selection for distinguishing single versus cross-talk activation pathways. **(a)** Illustration of two-state model fitting to nascent RNA mean and Fano factor data generated by the cross-talk pathway model. To match the rapid overshoot in nascent RNA dynamics, the two-state model requires an extremely large estimated *k*_on_; this, however, results in significant deviations in Fano factor fitting. **(b)** Accuracy of distinguishing single-pathway (two-state and three-state models) and cross-talk pathway models using nascent RNA moment data, across 100 datasets, 4 sample sizes (*N* = 10^2^, 10^3^, 5 × 10^3^, 10^4^), and 5 time point numbers (*n* = 2, 4, 8, 12, 24). Over 75% of datasets are correctly classified, with robust performance even for small *N* and *n*. **(c)** Fitting results of the two-state model to mRNA mean and Fano factor data from the cross-talk pathway model. Unlike nascent RNA (which shows large deviations), mRNA data exhibits only minor Fano factor discrepancies. **(d)** Accuracy of model selection using mRNA moment data under the same conditions as (b). While accuracy exceeds 75% for single-pathway datasets, it falls below 35% for cross-talk pathway datasets, indicating poor performance for distinguishing cross-talk pathways in mRNA data.

**Figure 6:**
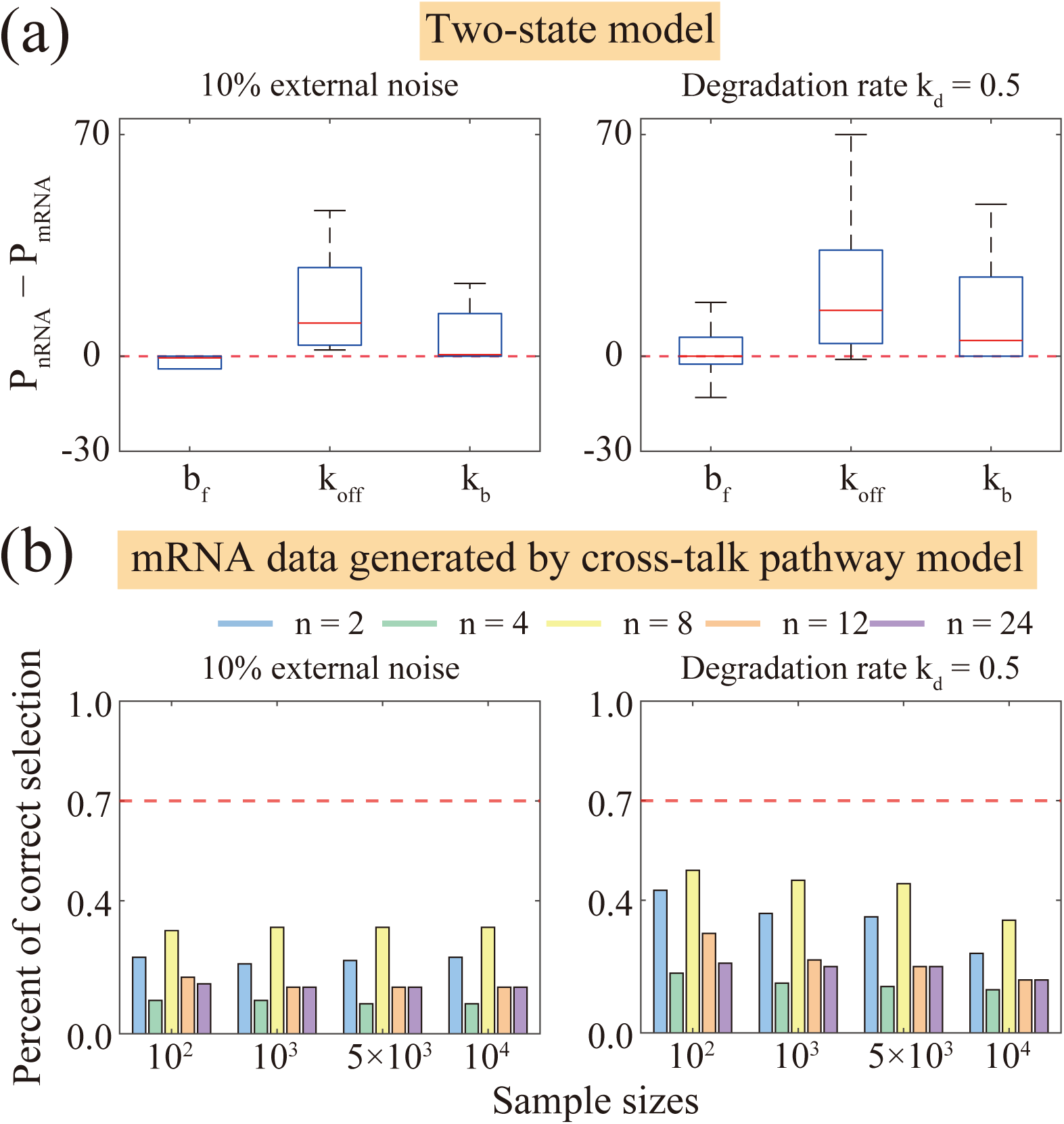
Impact of extrinsic noise and long mRNA lifetime on parameter estimation and model selection , across 100 datasets, 4 sample sizes (*N* = 10^2^, 10^3^, 5 × 10^3^, 10^4^), and 5 time points (*n* = 2, 4, 8, 12, 24). **(a)** Differences in accurate estimation percentages (*P*_nascentRNA_ − *P*_mRNA_) for parameters (*b_f_ , k*_off_*, k*_b_) under extrinsic noise (mRNA synthesis rate *k*_b_ with 10% log-normal noise) or long mRNA lifetime (degradation rate *k*_d_ = 0.5). Positive values indicate that nascent RNA outperforms mRNA. nascent RNA remains superior in estimating *k*_off_ and *k*_b_. **(b)** Accuracy of distinguishing cross-talk pathways from single-pathway models using mRNA moment data. For cross-talk pathway datasets, accuracy remains below 35% regardless of extrinsic noise or long lifetime.

To validate our model selection procedure, we utilized 100 synthetic nascent RNA datasets generated by each of the two-state, three-state, and cross-talk pathway models under four sample sizes (*N* = 10^2^, 10^3^, 5 × 10^3^, 10^4^) and five time points (*n* = 2, 4, 8, 12, 24). As shown in Fig. 5b, our model selection procedure can correctly identify the underlying generative model for over 75% of the tested datasets. More importantly, the accuracy of our method remains robust even when both the cell sample size and the number of time points are small. This represents a significant improvement over existing model selection methods, which typically show reduced accuracy for small *N* -particularly when *N* ≤ 10^2^ [26, 31]. However, our procedure may not be applicable to mRNA data. As illustrated in Fig. 5c, the two-state model still provides a good fit to mRNA data generated by the cross-talk pathway model, with no obvious deviations in Fano factors between the empirical data and the fitting curves. We further applied the model selection procedure to 100 synthetic mRNA datasets generated by each of the three models under the same four sample sizes and five numbers of time points. As shown in Fig. 5d, while the selection accuracy exceeds 75% for datasets generated by the two-state or three-state model, it drops below 35% for those generated by the cross-talk pathway model. In summary, our moment dynamics-based model selection procedure performs reliably for nascent RNA data but may fail when applied to mRNA data.

### Verification of results under extrinsic noise and longer mRNA lifetime

We have demonstrated that using nascent RNA moment data exhibits superior performance compared with mRNA data in both parameter estimation and model selection. However, these discussions did not account for extrinsic noise during mRNA production [19, 29] or the relatively longer lifetime of mRNA compared to that of nascent RNA [4, 10]. To shed light on how extrinsic noise and long mRNA lifetime impact the conclusions of this study, we systematically incorporate them to generate our synthetic data and carry out inferences. To incorporate extrinsic noise, we followed the approach described in [19, 33] by introducing noise into the mRNA synthesis rate *k*_b_ of the models. Specifically, we redefined the mRNA synthesis rate *k*_b_ as a log-normally distributed random variable, where its mean remains equal to the original *k*_b_ value, and its standard deviation is set to 0.1 times the mean-corresponding to a 10% noise level. To simulate the long lifetime of mRNA, we set the mRNA degradation rate to *k*_d_ = 0.5, which corresponds to a lifetime of 1*/k*_d_ = 2, while maintaining the transcription time of nascent RNA at *T* = 1. Then we generated synthetic mRNA moment datasets for all three models under conditions of extrinsic noise or long mRNA lifetime, using the same 100 parameter sets, 4 cell samples, and 5 time points, as in the previous sections.

For parameter estimation, we tested synthetic mRNA data generated by the two-state model and calculated the percentages *P*_mRNA_ under extrinsic noise or long lifetime. We then referenced the percentages of accurate parameter estimations from nascent RNA data, *P*_nascentRNA_, shown in Fig. 4, and computed the differences *P*_nascentRNA_ − *P*_mRNA_ for each of the following parameters: *b_f_ , k*_off_ and *k*_b_; see Fig. 6a. Under extrinsic noise, mRNA data still outperforms nascent RNA data in estimating *b_f_*but performs worse in estimating *k*_off_ and *k*_b_. Notably, the overall values of *P*_nascentRNA_ − *P*_mRNA_ are slightly larger than the corresponding values in Fig. 4c, indicating that extrinsic noise in mRNA data does affect parameter estimation accuracy. Furthermore, under the condition of a long mRNA lifetime, most *P*_nascentRNA_ − *P*_mRNA_ values exceed 0-this clearly demonstrates that nascent RNA data performs significantly better than mRNA data when mRNA lifetime is relatively long. In summary, nascent RNA data consistently outperforms mRNA data in the presence of either extrinsic noise or when lifetime of mRNA is longer.

For model selection, we evaluated synthetic mRNA datasets generated by all three candidate models under conditions of extrinsic noise or long lifetime. When mRNA data were simulated using either the two-state model or the three-state model, our fitting-based selection procedure demonstrated robust performance: it correctly identified the underlying generative model in over 75% of test datasets across all evaluated sample sizes and time points (Supplementary Fig. S4). In contrast, our procedure showed limited reliability in selecting the correct model when analyzing mRNA data generated by the cross-talk pathway model, regardless of whether extrinsic noise was introduced or lifetime was prolonged (Fig. 6b). Notably, extrinsic noise did not exert a significant impact on the overall accuracy of our model selection framework-a finding that highlights the algorithm’s resilience to stochastic perturbations in mRNA measurements.

### Verification of results using experimental transcription data

We validated procedure of our model selection (Single vs. multiple pathways) using real datasets: nascent RNA data of *CTT1* (a hyperosmotic stress-responsive gene in *S. cerevisiae*) [29], and mRNA data of the inducible P_lac*/*ara_ promoter in living *E. coli* cells under growth stimulation [34]. Both systems are hypothesized to be regulated by cross-talk pathways. From a biological regulation standpoint, *CTT1* exhibits dual regulatory requirements inherent to stress-responsive genes [35]: first, a spontaneous basal pathway with weak activation operates under normal growth conditions; second, under osmotic stress, its transcription is robustly induced via the MAPK HOG pathway-an osmostress-specific signaling cascade with strong activation capacity. For the P_lac*/*ara_ promoter, induction by arabinose and IPTG relies on competitive regulation between the repressor LacI and activator AraC, both of which target the promoter [34]. Specifically, LacI dissociation from the promoter and AraC binding to the promoter represent two functionally distinct regulatory pathways. From a data fitting perspective, the cross-talk pathway model can capture the dynamic mRNA number distributions of both systems (at the cost of certain computational overhead) [26, 36]. In contrast, single-pathway models fail to replicate the temporal bimodal features of the observed mRNA distribution. This underscores the necessity of incorporating competitive cross-talk regulation to faithfully describe their transcriptional dynamics.

We then applied our model selection procedure described in Fig. 5 to fit the transcriptional data of the two genes separately. For the nascent RNA data of *CTT1*, the two-state model could well capture the characteristic overshoot of the nascent RNA dynamic mean, as illustrated in Fig. 7a. Here, we normalized the time scale by setting the peak time of the nascent RNA dynamic mean to *T* = 1, with all other time points adjusted accordingly. Consistent with our prediction (Fig. 5a), the Fano factor generated by the two-state model was extremely small, leading to a significant discrepancy from the experimental data with ARE of 10.6, which exceeds the predefined threshold of 1. Thus, our method selected the cross-talk pathway model to explain the nascent RNA data of *CTT1* transcription. In contrast, for the mRNA data of the P_lac*/*ara_ promoter, the overshoot of the mean was not prominent (time was the product of real time and the mRNA degradation rate *k*_d_ = 0.014 min^−1^). Consistent with our prediction (Fig. 5c), this lack of distinct dynamic features resulted in no obvious differences between the experimental mean/Fano factor data and the fitting curve of the two-state model, as shown in Fig. 7b. Consequently, our method was ineffective for inferring the regulatory model from this mRNA data.

**Figure 7:**
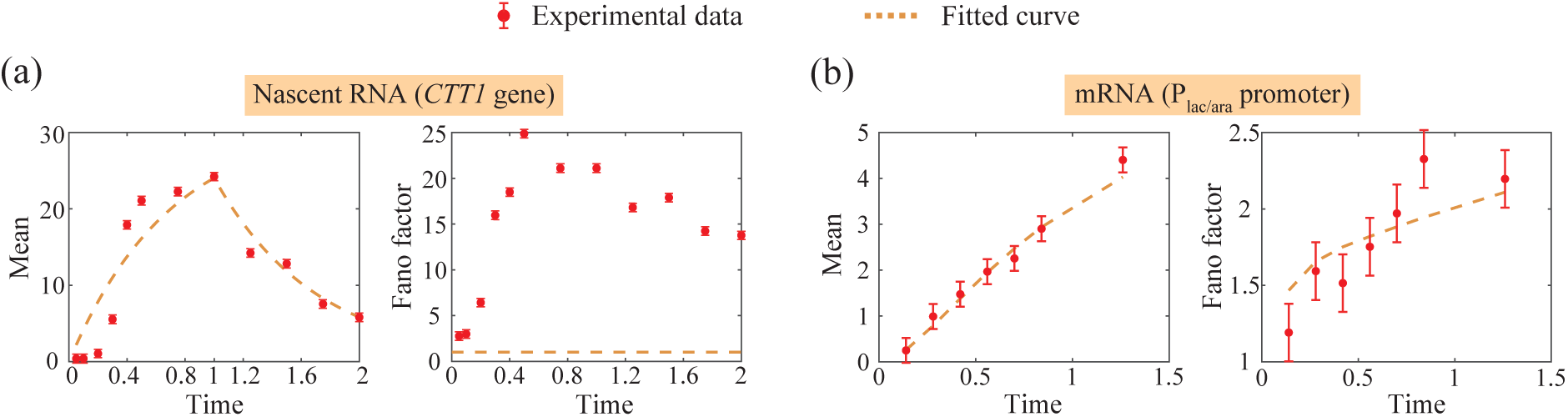
Two-state model fitting to *CTT1* nascent RNA (*S. cerevisiae*) and P_lac*/*ara_ promoter mRNA (*E. coli*) data [29, 34]. **(a)** For *CTT1*, the model captures nascent RNA mean overshoot but shows large Fano factor discrepancy (ARE = 10.6). **(b)** For P_lac*/*ara_, mRNA mean overshoot is indistinct, leading to no obvious fitting differences between experimental data and the model.

### Conclusions and discussions

Understanding how gene expression is regulated remains a central question in molecular biology. [1, 2, 4]. One of the most commonly used approaches to study the mechanisms of gene regulation is fitting theoretical models to mRNA copy number data. [26, 29, 31]. This approach is widely adopted because of its relative simplicity and the abundance of available mRNA copy number data. However, its effectiveness is severely limited when the cell sample size is small. This limitation poses a significant challenge, especially when dealing with experimental techniques such as single-molecule FISH, which typically have a restricted cell throughput [5]. To enhance the reliability of model inference based on mRNA data, it is essential to incorporate dynamical information, i.e., measurement of mRNA copy numbers at multiple time points [30]. However, fitting dynamical distribution data is computationally demanding, as it typically involves simulating chemical master equations and evaluating complex hypergeometric functions.

To address this computational challenge, an alternative approach is to perform model inference by fitting the moments of the mRNA distribution as a function of time. The advantage of moments-based approach is that both the moments can be expressed or approximated as elementary functions that are relatively easy to compute. This approach has successfully estimated parameters for the classical two-state model [31] and its extended auto-feedback variant [32]. However, mRNA production is inevitably accompanied by post-transcriptional noise, as mRNA molecules undergo various cytoplasmic decay pathways [29]; observed fluctuations in mRNA copy numbers may thus reflect not only the mechanisms of transcriptional regulation but also the stochasticity of post-transcriptional events. In contrast, nascent RNA, as an early intermediate in gene expression, may more accurately reflect the regulatory mechanisms at the level of transcription [19].

In this study, we developed a moments-based approach to infer models of transcriptional regulation using nascent RNA data, while systematically comparing its performance with that of mRNA data in both parameter estimation and model selection. We validated this approach using three models of transcriptional regulation: 1) two-state model, 2) three-state model, and 3) the cross-talk pathway model (Fig. 1). For these models, we derived analytical expressions for the transient and steady-state first two moments, as well as the Fano factor of the nascent RNA distribution (Tables 1-2). In particular, compared to mRNA , nascent RNA exhibits a more pronounced “overshoot” in the mean level as a function of time and a higher Fano factor (Fig. 2). This “overshoot” in mean is most significant for the cross-talk pathway model. Such a characteristic has been observed in the *HSP101* and *HsfA2* genes of Arabidopsis under heat induction [3]. These unique dynamic features of nascent RNA not only reflect its intrinsic advantage in preserving information about mechanisms of transcriptional regulation but also provide distinct discriminative signatures for model inference.

The principle of our moments-based approach involves simultaneously fitting mean *M* (*t*), second moment *S*(*t*) and Fano factor *η*(*t*) data by minimizing the corrected sum-of-squares difference *D_c_*(*θ*) as defined in Eq. 1. Intuitively, this approach might be expected to yield comparable to the traditional method of simultaneously fitting *M* (*t*) and *S*(*t*) [29, 31, 32]. This expectation holds true when fitting mRNA data but breaks down when applied to nascent RNA data (Fig. 3). The key reason lies in the unique “overshoot” dynamics of nascent RNA *M* (*t*): it forms a sharp peak within a narrow time window, a feature that is difficult to capture effectively through the fitting of *M* (*t*) and *S*(*t*) alone. Moreover, such fitting inaccuracies can be amplified in the discrepancy between the Fano factor data and its corresponding fitted curve. By incorporating *η*(*t*) into the fitting process, our approach forces the fitted curve to align more precisely with nascent RNA *M* (*t*) data within the peak region and the post-peak time window, which represents a key advantage of our method.

The superior parameter estimation performance of our approach over traditional least-squares methods stems from incorporating the Fano factor as an additional data dimension. This design complements existing multi-data integration strategies: for example, joint analysis of nuclear and cytoplasmic mRNA steady-state distributions has been shown to boost estimation accuracy and improve computational efficiency relative to single-compartment data [37]; similarly, integrating nascent RNA initiation time distributions with steady-state count data enables robust discrimination between regulatory mechanisms that static data alone cannot distinguish [38]. Our method complements these advances by focusing on time-dependent moments of nascent RNA, expanding multi-data integration to kinetic features, rather than solely static distributions. Notably, Bayesian methods-a prominent approach in transcriptional inference [15]-excel at quantifying parameter uncertainty and incorporating prior biological knowledge but suffer from high computational costs. In contrast, our approach relies on closed-form moment expressions, enabling efficient computation while preserving accuracy. This makes it a complementary tool-particularly valuable for high-throughput studies-while enriching the landscape of multi-data integration methods.

We applied this approach of parameter estimation to synthetic data with varying cell sample sizes and time points. Compared to the traditional method (fitting only *M* (*t*) and *S*(*t*)), our approach shows significantly better performance for nascent RNA data and marginally improved results for mRNA data (Fig. 3). Recent studies have suggested that for the two-state model, fitting synthetic steady-state nascent RNA distribution data with a large sample size of 10^4^ may yield more robust parameter estimation than fitting synthetic steady-state mRNA distribution data [19]. In contrast, we focus on the temporal as well as the steady-state moments of nascent RNA and mRNA (Fig. 4). In parameter estimation, nascent RNA data shows slightly lower reliability than mRNA data when estimating transcriptional frequency *b_f_* in the two-state and three-state models, but outperforms mRNA data in the cross-talk pathway model. In contrast, when estimating both the gene inactivation rate *k*_off_ and RNA synthesis rate *k*_b_, nascent RNA data consistently exhibits higher reliability than mRNA data. Notably, *k*_off_ is notoriously difficult to estimate accurately using both nascent RNA [19] and mRNA data [26, 33]. Even so, our proposed approach-when combined with nascent RNA data-still outperforms mRNA-based methods in estimating *k*_off_, thereby enhancing the overall estimation accuracy of parameters (*b_f_ , k*_off_*, k*_b_) (Fig. 4). For the two-state model, reliable estimation of all parameters using nascent RNA moment data requires either sample size *N* = 10^2^ with time point number *n* ≥ 4, or *N* ≥ 10^3^ with *n* ≥ 2. For the three-state model, reliable estimation demands at least *N* = 10^3^ with *n* ≥ 8, or *N* ≥ 5 × 10^3^ with *n* ≥ 4. However, for the cross-talk pathway model, tested *N* and *n* failed to achieve *>* 70% accuracy for *b_f_* and *k*_off_ estimation-consistent with prior findings that moments-based methods may struggle with in complex regulatory frameworks [29, 31].

The possible factor contributing to the cross-talk pathway model’s relatively inaccurate parameter estimation lies in the inherent definition of its pathway selection probabilities *q*_1_ (or *q*_2_). Unlike other kinetic parameters in the model, *q*_1_ and *q*_2_ are dimensionless-describing transcription factors (TFs) binding preferences to the promoter from the two competing pathways-rather than rates tied to specific biochemical steps. Biologically, TFs binding to promoter motifs typically involves at least one rate-limiting step (e.g., TF conformational changes or promoter chromatin remodeling), processes that would ideally be quantified by kinetic rates. Yet in the current cross-talk model, we assumed instantaneous TFs-promoter binding to highlight the core competitive cross-talk mechanism between the two pathways. This biologically motivated simplification omits critical kinetic details implicitly embedded in *q*_1_ and *q*_2_, introducing ambiguity into the inference of the other parameters in the model. Future work could explore more mechanistically detailed models-explicitly describing pathway selection’s biochemical processes (e.g., adding TFs-promoter binding/unbinding rates)-to assess how such refinements improve parameter estimation reliability for cross-talk systems.

We further applied our moments-based approach with nascent RNA data to model selection: distinguishing single activation pathway models (two-state or three-state model) from the cross-talk pathway model (Fig. 5). This relies on a key feature: the cross-talk pathway model generates a far higher mean peak and larger Fano factor than single-pathway models (Fig. 2). For the selection process, if nascent RNA data come from the cross-talk pathway model, fitting with a two-state model forces a very large gene activation rate *k*_on_ to match the *M* (*t*) peak. Yet this large *k*_on_ yields a low Fano factor in the fitted curve, leading to a mismatch between the Fano factor as predicted from the model and synthetic data. A predefined threshold check on this mismatch then identifies the model. Notably, this procedure achieves *>* 75% correct model identification across all tested sample sizes and time points. Importantly, it performs robustly even under small-scale experimental conditions with as few as only *N* = 100 and *n* = 2. This is a vital advantage for low-throughput experiments where sample and time resources are limited. In contrast, mRNA data fails to distinguish cross-talk pathway model data, with accuracy dropping to *<* 35%, highlighting our approach’s strength in leveraging nascent RNA’s unique dynamics.

In summary, this study proposes a moments-based approach that leverages nascent RNA’s unique dynamics in mean and Fano factor, establishing it as a more informative proxy than mRNA data for inferring regulatory mechanisms-including both parameter estimation and model selection. Notably, these conclusions are robust: we validated their reliability by introducing extrinsic noise during mRNA production and prolonging mRNA lifetime, with no significant decline in performance observed (Fig. 6). Despite these advances, limitations remain. First, experimental validation is needed to confirm *in vivo* measurability of nascent RNA’s dynamical features; recent single-cell sequencing technologies [3, 4] enable time-resolved nascent RNA profiling, offering an opportunity for empirical testing. Second, parameter estimation for complex models remains challenging: even with nascent RNA data, reliable estimation of the cross-talk pathway model fails to achieve accuracy even under large samples *N* = 10^4^ and time point numbers *n* = 24. Meanwhile, our model will grow more complex as we account for detailed RNAP premature detachment, pausing, and termination along the gene [39]. Future work could explore more efficient methods by combining nascent RNA and mRNA distribution data [29]. Third, our model selection cannot distinguish the two-state from the three-state model, likely due to similar activation regulation. Future studies may test additional nascent RNA data types-such as distributions of initiation time [40] and distributions of waiting time [41] between successive nascent RNA production events-to discriminate these models.

### Data accessibility

The MATLAB code for SSA, moment-based parameter inference and model selection across the three models (both nascent RNA and mRNA) is available at https://github.com/jfandwkw/jf-and-wkw

## Author Contributions

S.C. and F.J. conceived the original study. K.W. performed theoretical derivations. K.W., Y.L. and J.W. performed numerical simulations and data analysis. K.W., Y.L., S.C. and F.J. interpreted the theoretical results. S.C. and F.J. jointly wrote the manuscript with assistance from the co-authors.

## Acknowledgments

This work was supported by grants from the Natural Science Foundation of China (No. 12271118), Guangzhou Municipal Science and Technology Plan Basic and Applied Basic Research (No. 2025A03J3089).

## Appendix Tables

**Table 3:** Exact formulas for the coefficients of the three-state model (presented in Table 2 of the main text).

|  |  |
| --- | --- |
| $a_0$ | $\frac{k_{\text{on},1}k_{\text{on},2}k_b}{\alpha\beta k_d}$ |
| $a_\alpha$ | $-\frac{k_{\text{on},1}k_{\text{on},2}k_b}{\alpha(\beta-\alpha)(k_d-\alpha)}$ |
| $a_\beta$ | $-\frac{k_{\text{on},1}k_{\text{on},2}k_b}{\beta(\alpha-\beta)(k_d-\beta)}$ |
| $a_{k_d}$ | $-\frac{k_{\text{on},1}k_{\text{on},2}k_b}{k_d(\alpha-k_d)(\beta-k_d)}$ |
| $b_0$ | $\frac{k_b k_d (k_d + \alpha)(k_d + \beta) + k_b^2 (k_d + k_{\text{on},1})(k_d + k_{\text{on},2})}{\alpha\beta k_d^2 (k_d + \alpha)(k_d + \beta)} \cdot k_{\text{on},1} k_{\text{on},2}$ |
| $b_\alpha$ | $-\frac{k_b k_d (2k_d - \alpha)(k_d + \beta - \alpha) + 2k_b^2 (k_d + k_{\text{on},1} - \alpha)(k_d + k_{\text{on},2} - \alpha)}{\alpha k_d (\beta - \alpha)(k_d - \alpha)(2k_d - \alpha)(k_d + \beta - \alpha)} \cdot k_{\text{on},1} k_{\text{on},2}$ |
| $b_\beta$ | $-\frac{k_b k_d (2k_d - \beta)(k_d + \alpha - \beta) + 2k_b^2 (k_d + k_{\text{on},1} - \beta)(k_d + k_{\text{on},2} - \beta)}{\beta k_d (\alpha - \beta)(k_d - \beta)(2k_d - \beta)(k_d + \alpha - \beta)} \cdot k_{\text{on},1} k_{\text{on},2}$ |
| $b_{k_d}$ | $-\frac{k_b k_d \alpha \beta + 2k_b^2 k_{\text{on},1} k_{\text{on},2}}{\alpha\beta k_d^2 (\alpha - k_d)(\beta - k_d)} \cdot k_{\text{on},1} k_{\text{on},2}$ |
| $b_{2k_d}$ | $\frac{k_b^2 (k_{\text{on},1} - k_d)(k_{\text{on},2} - k_d)k_{\text{on},1} k_{\text{on},2}}{k_d^2 (\alpha - 2k_d)(\beta - 2k_d)(\alpha - k_d)(\beta - k_d)}$ |
| $b_{k_d, \alpha}$ | $-\frac{2k_b^2 (k_{\text{on},1} - \alpha)(k_{\text{on},2} - \alpha)k_{\text{on},1} k_{\text{on},2}}{k_d \alpha (k_d + \alpha)(\beta - k_d - \alpha)(k_d - \alpha)(\beta - \alpha)}$ |
| $b_{k_d, \beta}$ | $-\frac{2k_b^2 (k_{\text{on},1} - \beta)(k_{\text{on},2} - \beta)k_{\text{on},1} k_{\text{on},2}}{k_d \beta (k_d + \beta)(\alpha - k_d - \beta)(k_d - \beta)(\alpha - \beta)}$ |

**Table 4:** Exact formulas for the coefficients of the cross-talk pathway model (presented in Table 2 of the main text).

|  |  |
| --- | --- |
| $a_0$ | $\frac{k_{\text{on},1}k_{\text{on},2}k_b}{\alpha\beta k_d}$ |
| $a_\alpha$ | $-\frac{k_{\text{on},1}k_{\text{on},2}k_b - \alpha k_b(q_1k_{\text{on},1} + q_2k_{\text{on},2})}{\alpha(\beta - \alpha)(k_d - \alpha)}$ |
| $a_\beta$ | $-\frac{k_{\text{on},1}k_{\text{on},2}k_b - \beta k_b(q_1k_{\text{on},1} + q_2k_{\text{on},2})}{\beta(\alpha - \beta)(k_d - \beta)}$ |
| $a_{k_d}$ | $-\frac{k_{\text{on},1}k_{\text{on},2}k_b - k_d k_b(q_1k_{\text{on},1} + q_2k_{\text{on},2})}{k_d(\alpha - k_d)(\beta - k_d)}$ |
| $b_0$ | $\frac{k_b k_d(k_d + \alpha)(k_d + \beta) + k_b^2(k_d + k_{\text{on},1})(k_d + k_{\text{on},2})}{\alpha\beta k_d^2(k_d + \alpha)(k_d + \beta)} \cdot k_{\text{on},1}k_{\text{on},2}$ |
| $b_\alpha$ | $-\frac{k_b k_d(2k_d - \alpha)(k_d + \beta - \alpha) + 2k_b^2(k_d + k_{\text{on},1} - \alpha)(k_d + k_{\text{on},2} - \alpha)}{\alpha k_d(\beta - \alpha)(k_d - \alpha)(2k_d - \alpha)(k_d + \beta - \alpha)} \cdot [k_{\text{on},1}k_{\text{on},2} - \alpha(q_1k_{\text{on},1} + q_2k_{\text{on},2})]$ |
| $b_\beta$ | $-\frac{k_b k_d(2k_d - \beta)(k_d + \alpha - \beta) + 2k_b^2(k_d + k_{\text{on},1} - \beta)(k_d + k_{\text{on},2} - \beta)}{\beta k_d(\alpha - \beta)(k_d - \beta)(2k_d - \beta)(k_d + \alpha - \beta)} \cdot [k_{\text{on},1}k_{\text{on},2} - \beta(q_1k_{\text{on},1} + q_2k_{\text{on},2})]$ |
| $b_{k_d}$ | $-\frac{k_b k_d \alpha \beta + 2k_b^2 k_{\text{on},1} k_{\text{on},2}}{\alpha\beta k_d^2(\alpha - k_d)(\beta - k_d)} \cdot [k_{\text{on},1}k_{\text{on},2} - k_d(q_1k_{\text{on},1} + q_2k_{\text{on},2})]$ |
| $b_{2k_d}$ | $\frac{k_b^2(k_{\text{on},1} - k_d)(k_{\text{on},2} - k_d)}{k_d^2(\alpha - 2k_d)(\beta - 2k_d)(\alpha - k_d)(\beta - k_d)} \cdot [k_{\text{on},1}k_{\text{on},2} - 2k_d(q_1k_{\text{on},1} + q_2k_{\text{on},2})]$ |
| $b_{k_d,\alpha}$ | $-\frac{2k_b^2(k_{\text{on},1} - \alpha)(k_{\text{on},2} - \alpha)}{k_d \alpha(k_d + \alpha)(\beta - k_d - \alpha)(k_d - \alpha)(\beta - \alpha)} \cdot [k_{\text{on},1}k_{\text{on},2} - (k_d + \alpha)(q_1k_{\text{on},1} + q_2k_{\text{on},2})]$ |
| $b_{k_d,\beta}$ | $-\frac{2k_b^2(k_{\text{on},1} - \beta)(k_{\text{on},2} - \beta)}{k_d \beta(k_d + \beta)(\alpha - k_d - \beta)(k_d - \beta)(\alpha - \beta)} \cdot [k_{\text{on},1}k_{\text{on},2} - (k_d + \beta)(q_1k_{\text{on},1} + q_2k_{\text{on},2})]$ |

